# Dissecting the factors shaping fish skin microbiomes in a heterogeneous inland water system

**DOI:** 10.1101/739748

**Authors:** Yaron Krotman, Timur M. Yergaliyev, Rivka Shani Alexander, Yosef Avrahami, Amir Szitenberg

**Author notes:** Equal contributors (Yaron Krotman, Timur M. Yergaliyev).

## Abstract

Fish skin microbiomes are rarely studied in inland water systems, in spite of their importance for fish health and ecology. This is mainly because fish species distribution often covaries with other biotic and abiotic factors, complicating the study-design. We tackled this issue in the northern part of the Jordan River system, in which a few fish species geographically overlap, across a steep gradients of water temperature and salinity. Using 16S rRNA metabarcoding, we studied the water properties that shape the skin bacterial communities, and their interaction with fish taxonomy. We found that considering the skin-community contamination by water microbial community is important, even when the water and skin communities are apparently different. With this in mind, we found alpha diversity of the skin-communities to be stable across sites, but higher in bentic loaches, compared to other fish. Beta diversity was found to be different among sites and to weakly covary with the dissolved oxygen, when treated skin-communities were considered. In contrast, water temperature and conductivity were strong factors explaining beta diversity in the untreated skin-communities. Beta diversity differences between co-occurring fish species emerged only for the treated skin-communities. Metagenomics predictions highlighted the microbiome functional implications of excluding the water-communities contamination from the fish skin-communities. Finally, we found that human induced eutrophication promotes dysbiosis of the fish skin-community, with signatures relating to fish health. This finding was in line with recent studies, showing that biofilms capture sporadic pollution events, undetectable by interspersed water monitoring.

## Introduction

The importance of the cutaneous mucus in fish is well established; The teleost epidermal mucus provides mechanical protection against physical and biological harm thanks to its viscosity and high turnover [1,2], and it contains agents taking part in ecological interactions [3]. Additionally, it is a primary immune response site, in which the innate immune system and antimicrobial peptides are highly active [4]. Other biochemical activities involving defensins, lysozymes and lectin-like agglutinins additionally respond to pathogens [5]. In contrast, many mutualistic and commensal microbes are well adapted to use the mucus as adhesion site and can evade the defence mechanisms it provides [6]. This community also interferes with infections [7–9], *via* competition or antagonistic interactions [10,11]. Dysbiosis of the skin microbial community can drive it out of homoeostasis and promote infection [12], although not every perturbation in the microbiome must lead to the loss of function [13].

Although the skin microbiome in fish has not been the focus of microbiome research, some important progress has been made by a few research groups. The skin microbiome is known to be affected by both environmental and fish-species dependant factors [14,15], with evidence for co-phylogeny in coral reef fish [15]. On the population level, however, the existence of microbiome covariation with host genetics is inconsistent among systems [16–18]. Interpopulation variation appears to rely, in part, on variable resolution of antagonistic relationships among microbial species [17]. Capture stress has been shown to correlate with microbiome contamination, in particular by *Vibrio* spp. [19]. Conversely, perceived opportunistic pathogens such as *Vibrio* spp. appear to constitute small fractions of normal microbiomes and culture dependent techniques grossly over-represent them [20]. Additional studies identified stress indicators [21] and probiotics candidates [22], both with conceivable applications in aquaculture and nature conservation, as well as the finding that captivity reduces the skin microbiome biodiversity [18,23,24]. Consistent salinity bioindicators were also recovered in an experimental system utilizing euryhaline fish [25].

While most of the current research is targeted at fish species with commercial relevance [2,26–29] or food safety [30], a few studies have dissected wild fish communities or populations, utilizing deep-sequencing culture-independent methods [15,16,31] and leaving the vast majority of wild habitats unexamined [32]. In the wild, particularly in fragmented and heterogeneous inland water systems, it is difficult to test the effect of geographically varying abiotic conditions on a given species, since the fish community composition often covaries with them [25].

In this work, we have sampled the upper reaches of the Jordan River system and Springs Valley streams, north and south of the Sea of Galilee, respectively. This range includes heterogeneous sites, differing in fish community composition and water properties. We sampled mostly in nature reserves, although three of the sites suffer human-induced eutrophication, one of which is a settling pool, and two others receiving fish-farm and fish-pond outlets. The geographic range of a few fish species in this part of the system partially overlap, thus allowing us to study host and site dependent effects on fish skin microbiomes. Due to the sensitivity of the sampled ecosystem, we employed a non-destructive sampling procedure, swabbing the captured fish on site and immediately releasing them. Our results reveal effects of both fish-species and sampling-site on the skin microbiome, highlight the importance of considering the background microbial contamination of the swab samples by the water, and show that eutrophication may drive the skin microbiome to dysbiosis.

## Results

### Sampling

To study the microbial diversity in freshwater fish skin and the factors shaping it, we have sampled a cumulative number of 14 species from 17 locations representing three streams north of the Sea of Galilee (three to six sites in each stream), and two streams to its south (one and two sites per stream). We will hereafter denote the two regions the “northern” and “southern” basins (Fig. 1; Table S1). Additionally, we collected two liters of water in each site. In total, we accumulated 176 fish-skin swab samples and 17 water bottles. In the northern basin *Capoeta damascina* (Cyprinidae) were collected from all sites in the Hermon (H) and Snir (S) streams, and from two sites in the Jordan River (J). The species most co-occurring with C. *damascina* was *Oxynoemacheilus insignis* (Nemacheilidae), which was found in three H sites, one S site and one J site. Unlike *C. damascina, O. insignis* was also captured in Tel-Saharonim Stream (T, southern basin). Another relatively widely dispersed group included the Tilapiine (Cichlidae) species *Coptodon zillii* (formerly *Tilapia*), *Sarotherodon galilaeus* and hybrids of *Oreochromis aureus*, which were found in three J sites in the northern basin, co-occurring with *C. damascina* in one site, and in the two southern basin streams, co-occurring with *O. insignis* in one site. The remaining species, belonging to Cyprinidae, Haplochrominae (Cichlidae), Poeciliidae and Mugilidae, had a narrow geographic rage and a small geographic overlap with other species (Table S1).

**Fig. 1:**
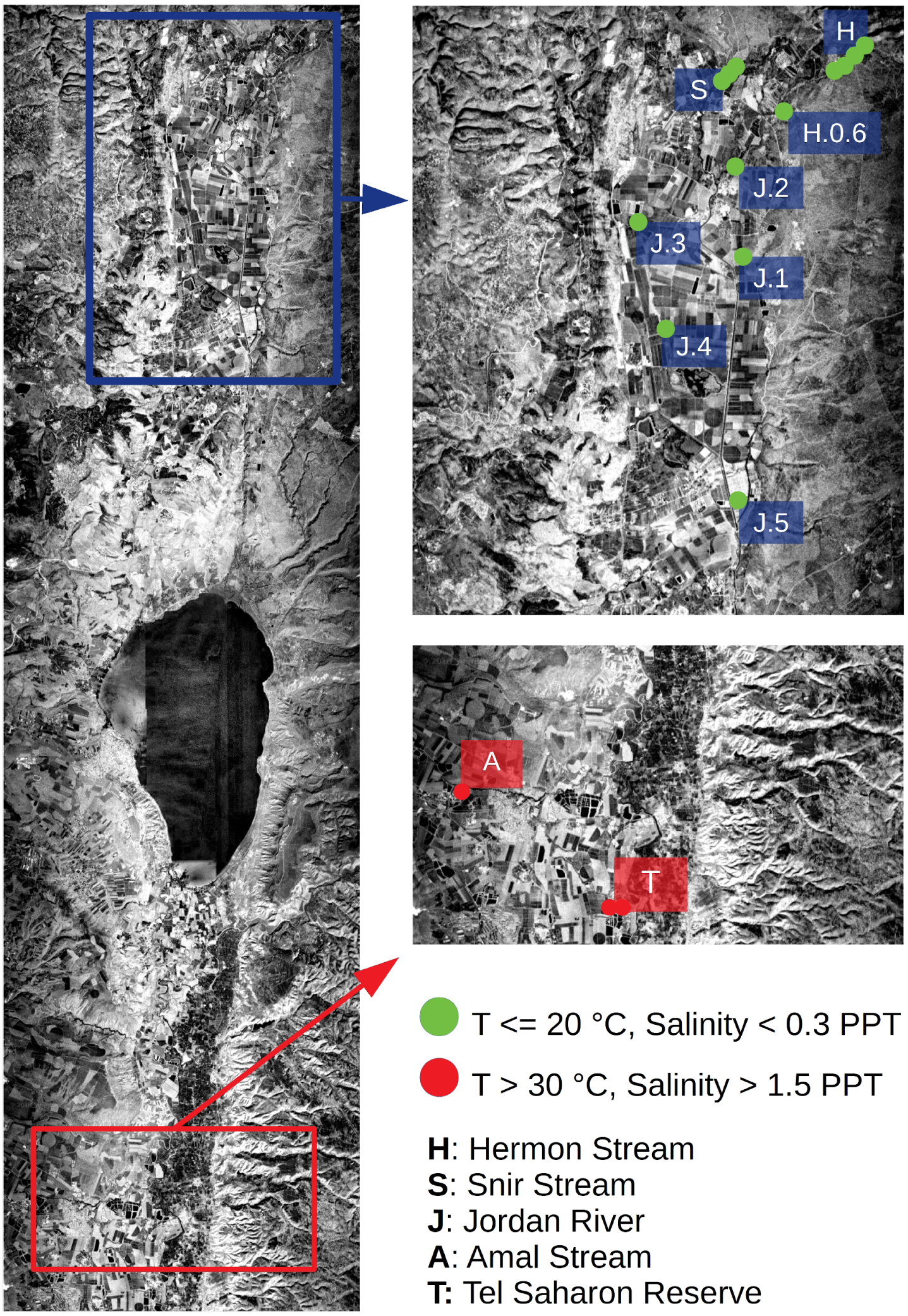
Sampling area. Samples were collected north (green bullets) and south (red bullets) to the Sea of Galilee, across temperature and salinity gradients.

### Sequence data: “raw-swab” and “skin-corrected” bacterial communities

To minimize our impact on the sampling site, we rubbed fish along their lateral line on site, using sterile swabs, and immediately released them. This method resulted in variable DNA quantities retrieved from each swab, and a subset of samples was selected post-hoc. According to alpha diversity rarefaction curves, the alpha diversity in both swab and water samples was thoroughly represented by 1000 sequences or more (Fig. S1). Consequently, after exclusion of organelle reads that may amount to as much as half the sequence data, we retained 120 fish-skin samples and 10 water samples, with a mean sequence-read count of 2996 sequence reads, ranging from 1022 and 6686 reads individual samples. We considered this dataset to represent “raw swab communities”. We further filtered the biom table to include only amplicon sequence variants (ASV) that were unique to swab samples, or that had significantly higher relative abundance in swab samples than in water samples, based on Benjamini-Hochberg corrected [33] Mann-Whitney U-test [34]. This data set was denoted the “corrected skin communities”. Throughout the results, we address both the raw swab communities and the corrected skin communities to study the effect of this analytic procedure.

### Key bacterial amplicon sequence variants in the fish skin microbiome

The bacterial classes recovered from raw swab communities, having the highest median relative abundances (Fig. 2A, gray boxes), belonged to Alphaproteobacteria (12%), Actinobacteria (11%), Gammaproteobacteria (10%), Bacilli (3%) and Fusobacteriia (3%). The corrected skin community (Fig. 2A, orange boxes), had higher representation of Bacilli (7%) and Fusobacteriia (4%), and lower representation of Gammaproteobacteria (6%) and Alphaproteobacteria (8%), in comparison with the raw swab communities. It is noteworthy that although not very abundant in the raw or corrected communities, class Bacteroidia (Bacteroidetes) had a much higher median relative abundance in the raw swab communities (3%)than in the corrected skin communities (0%).Prominent genera (Fig.2B),mostly belonging to these classes, included *Cetobacterium* sp. (Fusobacteriia, 3% and 4% in the raw and corrected community respectively), *Anaerobacillus* sp. (Bacilli,1% and 2%) and *Skermanella* sp. (Alphaproteobacteria, 2% and 4.5%).

**Fig. 2:**
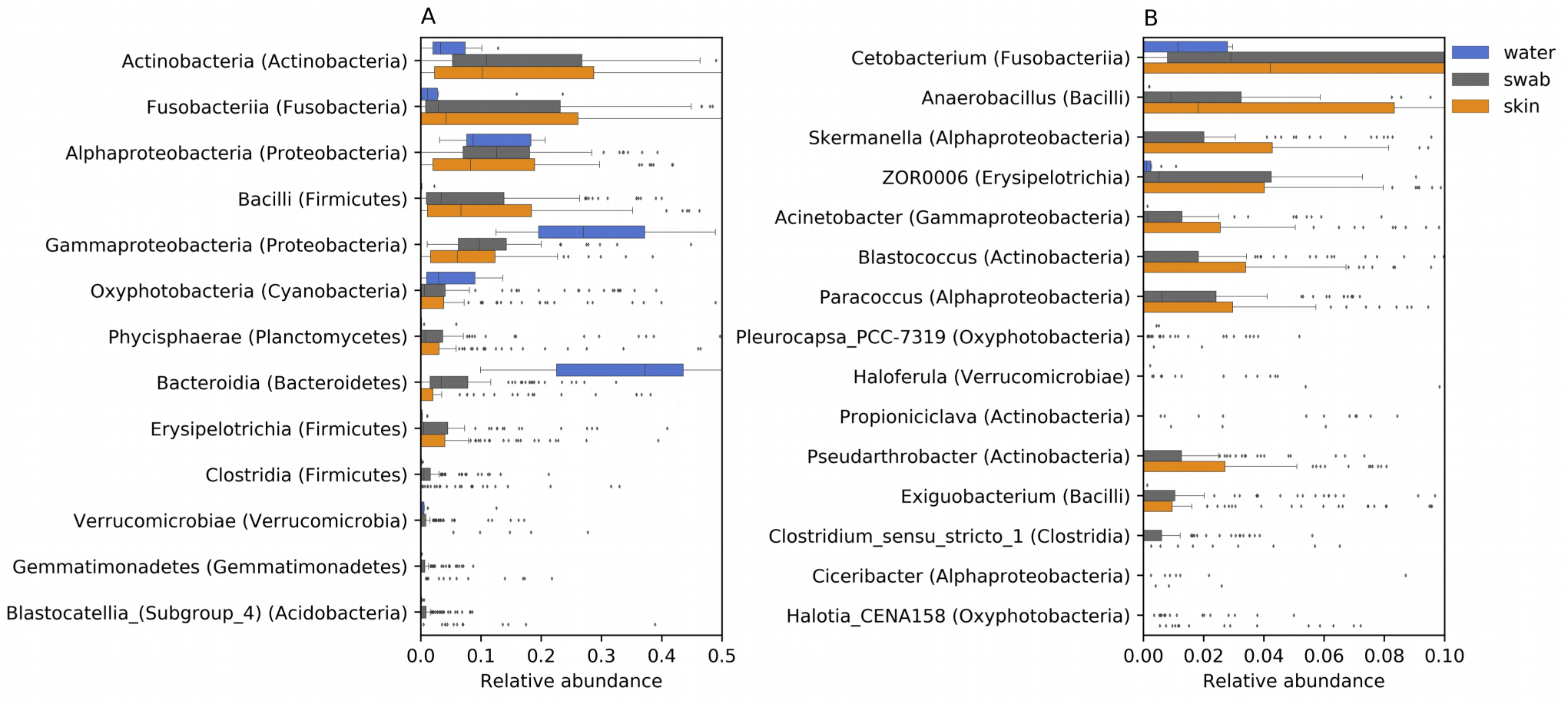
Relative abundance distributions of (A) bacterial classes (phyla) and (B) genera (classes) assignable to amplicon sequence variants (ASVs) recovered from water samples (blue boxes), skin samples (grey boxes) and corrected skin communities (orange boxes).

### Bacterial diversity

The following approach was taken to study the factors shaping the fish skin microbiome. Alpha and beta diversity were quantified with Faith’s phylogenetic diversity (Faith PD) index [35] and an unweighted-UNIFRAC [36] pairwise distance matrix, respectively. Significance differences among location and fish taxonomy categories were then tested with Benjamini-Hochberg corrected [33] Kruskal-Wallis [37] and PERMANOVA [38] tests, for alpha and beta diversity, respectively. Principal coordinates analyses (PCoA) [39,40] were used to visualize beta diversity clusters and the proportion of total variance they explain, coupled with biplot analyses [40], to detect the ASVs that change among the PCoA clusters. ANCOM tests [41] were used to identify ASVs that vary between sites or fish taxa. We further used Pearson correlation [42] to study the correlation between the water temperature, conductivity, pH or dissolved-oxygen, and the Faith PD values, or the first or second PCoA axis values. The entire procedure was carried out twice, for the raw swab communities and the corrected skin communities.

### Alpha diversity

Alpha diversity results are summarised in Fig. 3. Similar mean Faith PD values were found in raw swab communities (8.6 ± 2.6 SD) and water sample communities (9.2 ± 2.2), compared to the lower values computed for corrected skin communities (3.7 ± 0.8). When considering the raw swab communities, both the stream (p-value = 1.8×10^-11^) and fish family or tribe (p-value = 0.002) were significant factors, with many significant pairwise differences among streams (10^-7^ < q-value < 0.01; Fig. 3A; Table S2). Some additional significant differences were found among fish taxa, but only among largely non-overlapping fish families (Cyprinidae and Haplochrominae; q-value = 0.047, Cyprinidae and Tilapiinae; q-value = 0.047; Fig. 3B; Table S3). For these pairs of taxa, we cannot tease apart the location effect from that of fish taxonomy due to the covariance of the two factors. Temperature (Fig. 3C), conductivity (Fig. 3D) and pH (Fig. 3E), explained large proportions of the variance in raw swab community Faith PD values (R^2^= 0.37, 0.29 and 0.1, respectively).

**Fig. 3:**
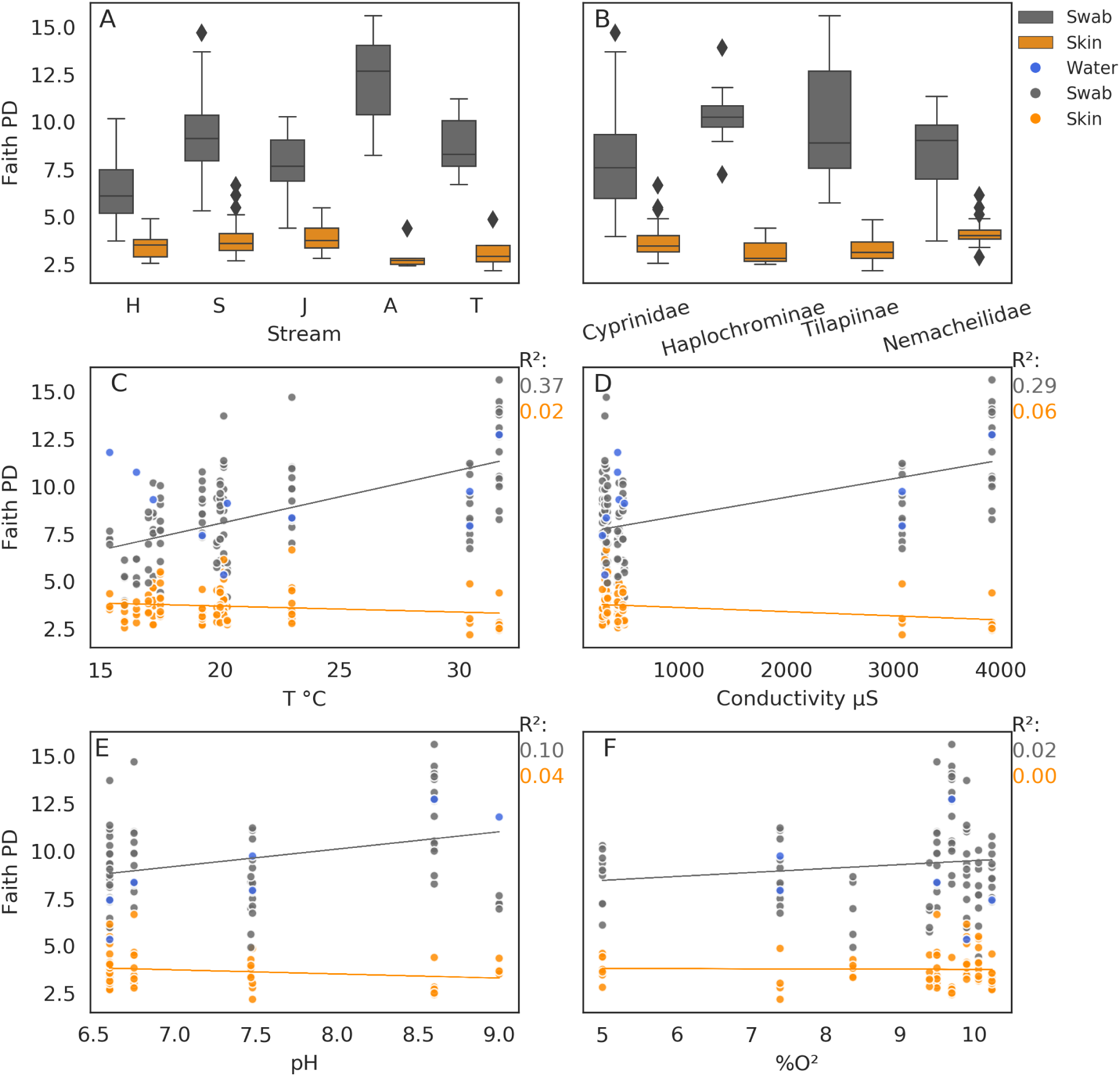
Bacterial alpha diversity. Faith PD distribution in swab bacterial communities (grey) and corrected skin communities (orange) in (A) streams and (B) fish families / tribes, and their correlations with water (C) temperature, (D) conductivity, (E) pH and (F) percent dissolved oxygen. Water sample Faith PD values in blue.

When considering the corrected skin communities, all pairwise differences between streams were non-significant, in contrast with the raw swab communities results. The only significant differences found were between Nemacheilidae and each of its co-occurring fish taxa Cyprinidae (q-value = 0.015) and Tilapiinae (q-value = 0.015). For the corrected skin communities, the water temperature, conductivity and salinity, no longer explained Faith PD values (R2= 0.02, 0.06 and 0.04, respectively). The Faith PD values of the water communities covaried with those of the raw swab communities, with the exception of sites with low water temperatures (Fig. 3C).

### Beta diversity

Both stream-based (Table S4) and fish-family based (Table S5) groupings were globally significant for the raw and corrected communities (p-value = 0.001), as well as pairwise stream comparisons (0.001 < q-value < 0.004). However, significant differences between pairs of fish taxa that co-occur geographically were found only for the corrected skin communities. These pairs included Nemacheilidae and its co-occurring fish taxa (Cyprinidae, Haplochrominae and Tilapiinae with q-value < 0.007 in the three comparisons). Differences between co-occurring fish taxa were not recovered for raw swab communities.

PCoA results for the raw swab communities, water sample communities and corrected skin communities are shown in Fig. 4A-B. For the raw and corrected communities, the first PCoA axis explained 14.7% and 14.8% of the total variance, respectively, and the second 7.2% and 10%, respectively. Despite the similar percent of explained variance between the raw and corrected communities, north and south basin distinctiveness was lost in the corrected dataset (Fig. 4B). Water sample communities from the northern basin clustered separately from raw-swab samples from the northern basin, but were similar to water samples and raw-swab samples from the southern basin (Fig. 4A).

**Fig. 4:**
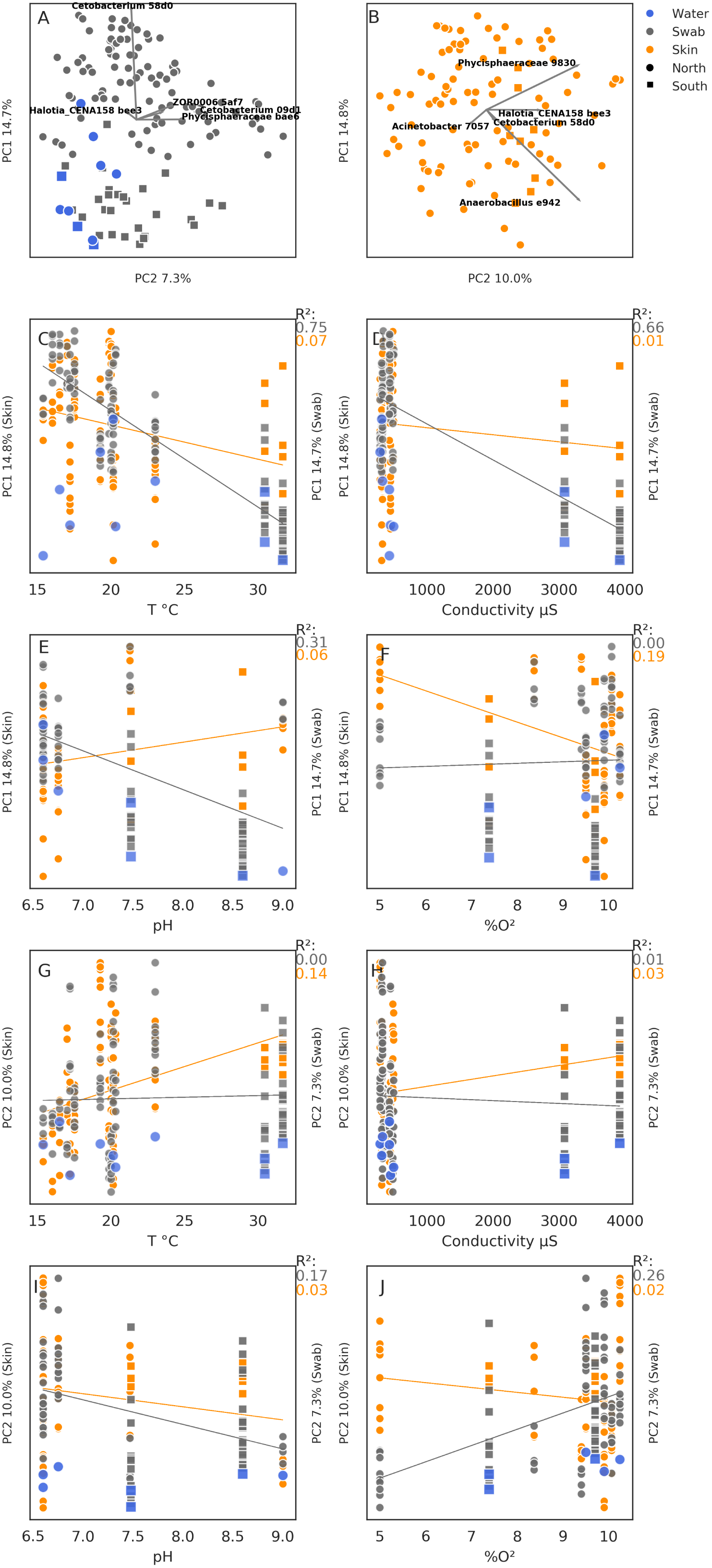
Beta diversity across the study area. Principal coordinates analysis (PCoA) of (A) swab bacterial communities and (B) corrected skin bacterial communities. The first and second PCoA axes correspond to the Y and X axes of each plot, respectively. The percent variance explained by each axis is denoted as the axis label. The four most important ASVs and their effect sizes are indicated by biplot arrows. Subplots C-J demonstrate the correlation of the first (C and F) and second (G and J) PCoA axis values with the temperature (C and G), Conductivity (D and H), pH (E and I) and dissolved oxygen (F and J). Circle and square markers denote the northern and southern basins respectively. Blue, grey and orange markers denote water, swab and corrected skin communities, respectively.

The explanatory ASVs changed between the raw and corrected communities, with *Cetobacterium* (ASV 58d0), a salinity bioindicator [25], having the strongest effect for the raw swab communities (Fig. 4A), and the anaerobes [43] Phycisphaeraceae (ASV 9830) and *Anaerobacillus* (ASV e942) for the corrected communities (Fig. 4B). Accordingly, In Fig. 4A, temperature, conductivity and pH (Fig. 4C-E) correlated strongly with the first PCoA axis values of the raw swab communities (R^2^= 0.75, 0.66 and 0.31, respectively), but this effect was mostly lost for the corrected skin community values (R^2^= 0.07, 0.01 and 0.06, respectively) and a weak correlation with the dissolved oxygen was observed instead (Fig. 4F, R^2^= 0.19). The second PCoA axis had weaker correlations with any of the water measurements than the first axis. For the second axis, the raw swab community values correlated with pH and dissolved oxygen measurements (R2= 0.17 and 0.26, respectively) and the corrected skin community values correlated with the temperature measurements (R^2^= 0.14).

To summarize, beta diversity in the raw swab communities is best explained by the water salinity or temperature. The corrected skin communities, however, are less affected by water characteristics, of which dissolved oxygen level is the strongest. Accordingly, in the raw swab communities, a salinity bioindicator bacterium varies the most, whereas for the corrected skin communities we detect large variations in anaerobic bacteria. It is important to note that dissolved oxygen measurements were not taken in the H stream, and thus the strength of this finding is tentative.

### Basin specific PCoA

To further investigate the relationship between the fish taxonomy and the skin microbiome, we carried out another PCoA, separating the northern and southern basins to increase the geographic range overlap of the included fish taxa in each analysis (Fig. 5). This analysis supported the importance of the sampling site in explaining the beta diversity in the raw swab communities (Fig. 5A and B, addressing the northern and southern basins, respectively, with marker shapes representing the different streams). However, stream separation was reduced when analysing the corrected skin community in the northern basin (Fig. 5C). This analysis further exposes a clear separation between Nemacheilidae and Cichlidae (Haplochrominae + Tilapiinae), for the raw swab communities (Fig. 5A) and more so for the corrected skin communities (Fig. 5C). According to ANCOM test, the bacterium explaining the difference between Nemacheilidae and Cichlidae is *Exiguobacterium* (ASV 0cb4) for both the raw and corrected communities.

**Fig. 5:**
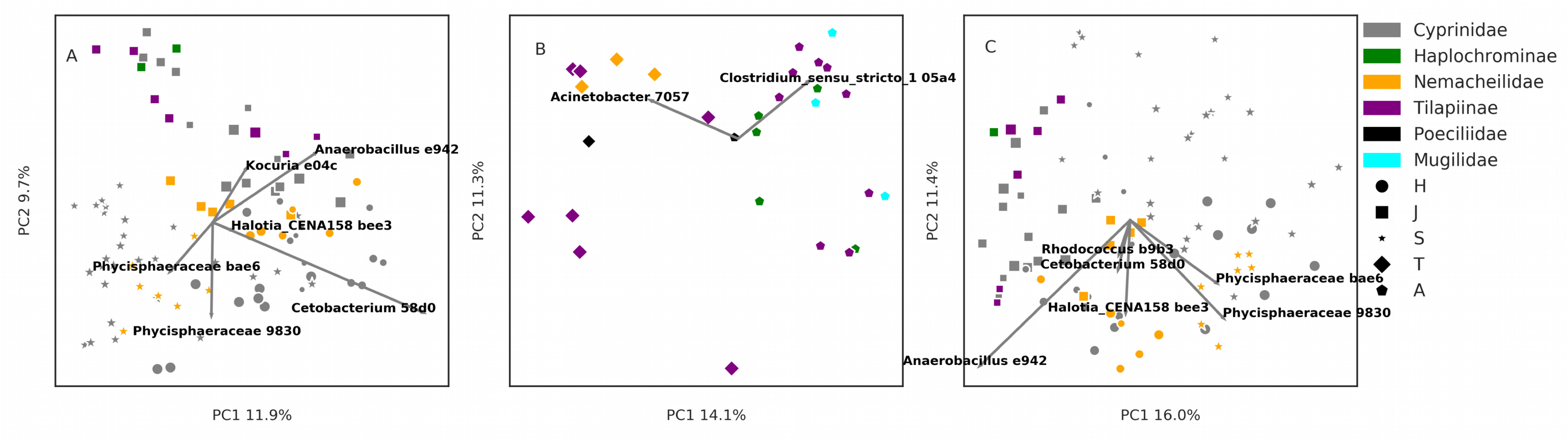
Beta diversity in the northern (A and C) and southern (B) basins. PCoA of (A and B) swab bacterial communities and (C) corrected skin bacterial communities. The first and second PCoA axes correspond to the X and Y axes of each plot, respectively. The percent variance explained by each axis is denoted as the axis label. The 2 - 4 most important ASVs and their effect sizes are indicated by biplot arrows.Marker colors denote different fish taxa, and marker shapes the different streams. Marker sizes denote specific sites within streams.

### Proteobacteria - Bacteroidetes ratios reveal dysbiosis in eutrophic sites

The ratio between Proteobacteria and Bacteroidetes is associated with fish health, with compromised individuals having increased Bacteroidetes relative abundances [44]. We compared the relative abundances of these phyla among the sampling sites to derive ecological insight (Fig. 6), checking both the raw swab communities (Fig. 6A) and the corrected skin communities (Fig. 6B). A clear connection emerged for the corrected skin community, associating elevated Bacteroidetes relative abundances and reduced Proteobacteria relative abundances with human induced eutrophication, site H.0.6 being a settling pool, site J.1 a fish pond outlet and sites S.2 and S.3 situated just upstream and downstream of a fish farm outlet. T.2, which was not initially categorised as eutrophic, also had a high relative abundance of Bacteroidetes. This is a shallow site amids agricultural land (Figure S2).

**Fig. 6:**
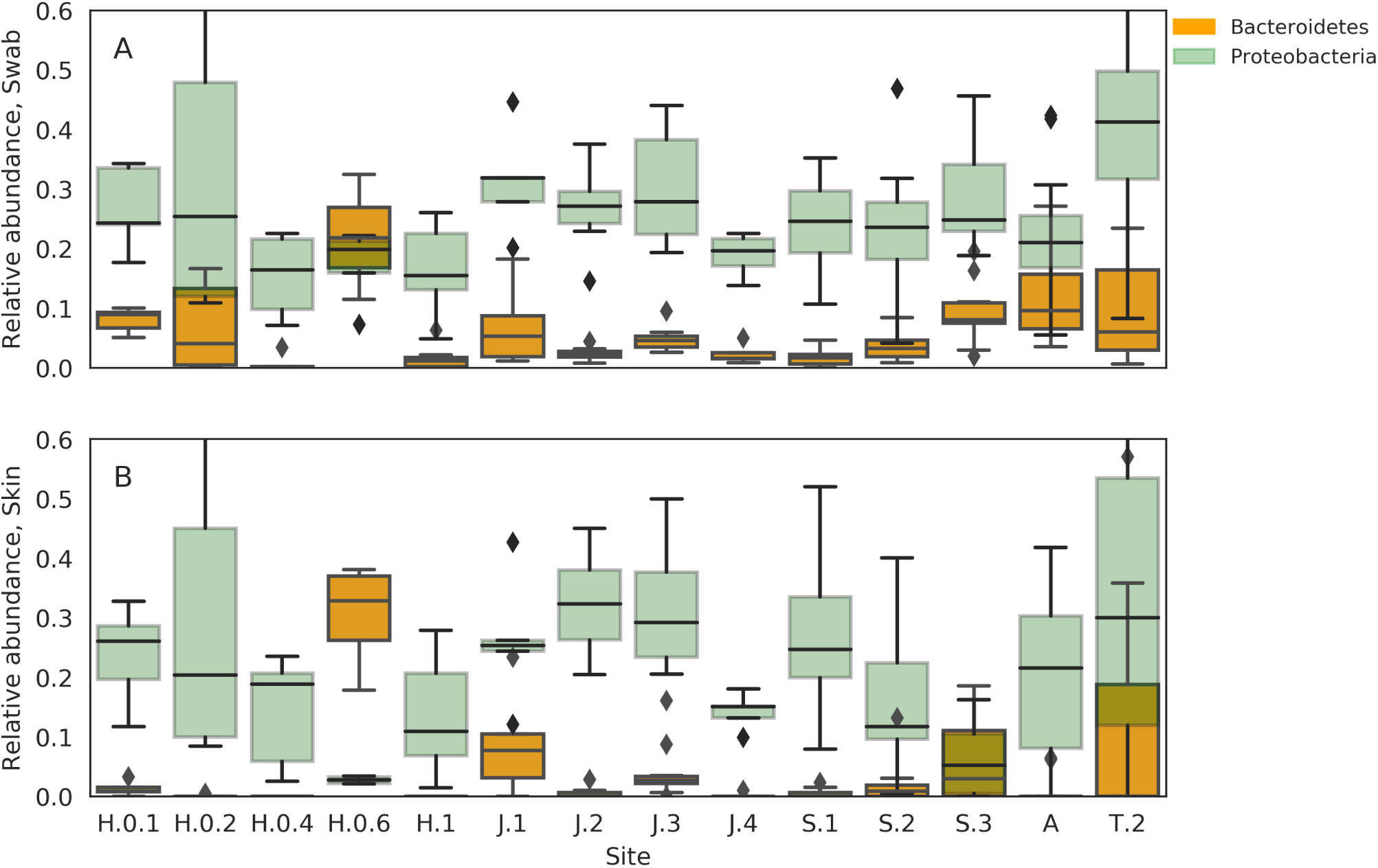
Relative abundances of Bacteroidetes (orange) and Proteobacteria (green) in the (A) raw swab bacterial communities and (B) corrected skin microbial communities.

### Predicted metabolic differences between the raw swab communities and the corrected skin communities

To understand the effect of water background contamination on the inference of fish skin microbiomes and the reconstruction of metabolic models that would be recovered from their metagenomes, we employed PICRUSt [45]. PICRUSt predicts a metagenome based on ASVs and bacterial genomes available in online databases. Fig. 7 summarizes the number of metabolic pathways with a significantly different relative abundance between the raw and corrected predicted metagenomes, according to their KEGG category. The most frequent KEGG categories with significantly different pathway representations were “Biosynthesis of secondary metabolites”, “Microbial metabolism in diverse environments” and “Biosynthesis of antibiotics”. This indicates that the raw swab communities and the corrected skin communities would produce different metabolic models, with respect to the ecological function of their members.

**Fig. 7:**
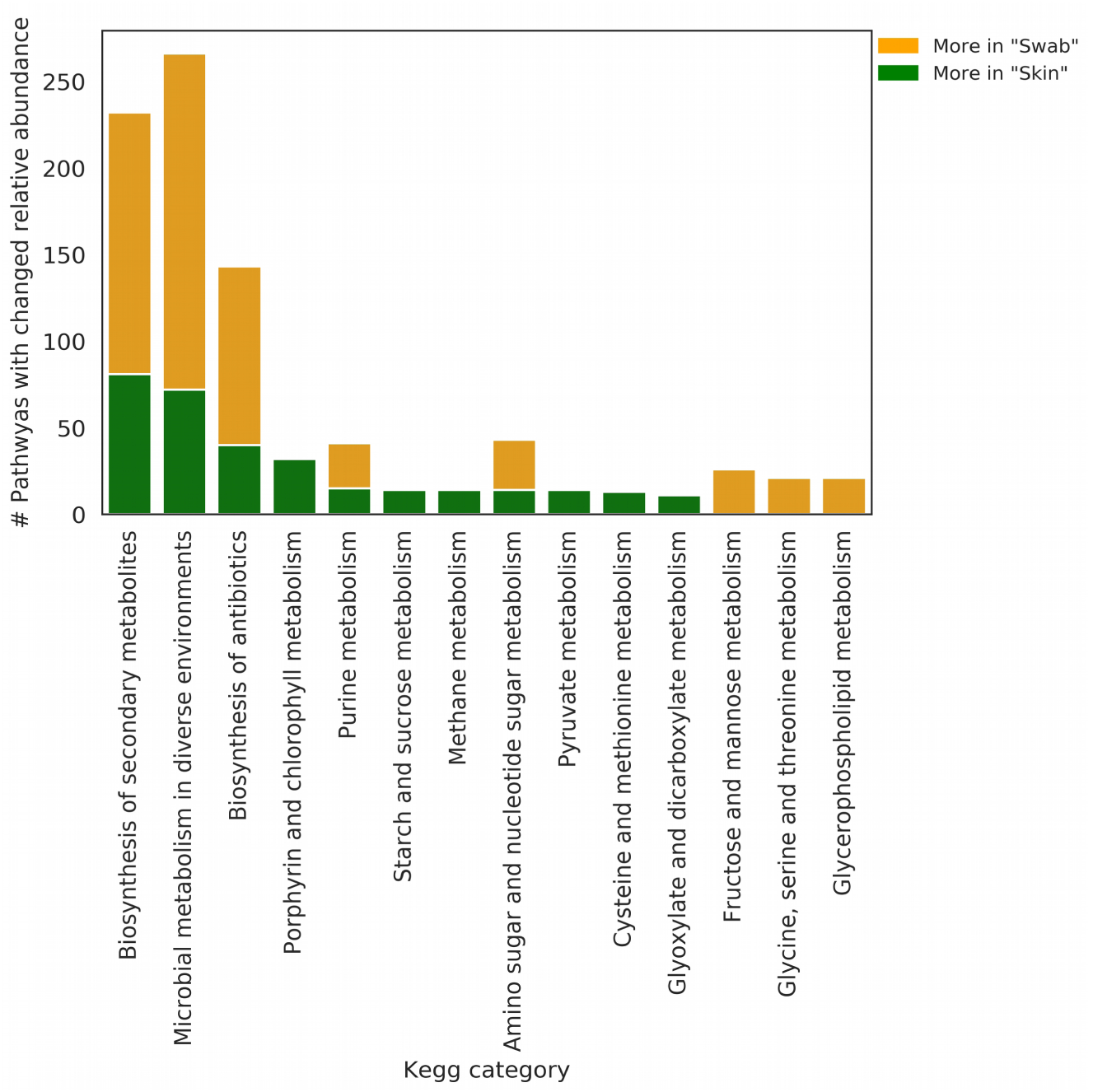
Functional differences between the raw swab bacterial communities (orange) and corrected skin communities (green). Each bar denotes the number of KEGG pathways with a significant relative abundance difference between the raw and corrected communities, in each of the KEGG pathway categories (x-axis).

## Discussion

### Freshwater fish-skin microbiome

Few studies have investigated skin microbiomes in freshwater fish, and it is not clear if it is fundamentally different than those of marine fish. Larsen et al. [14] have sampled the catadromous *Mugil cephalus* in marine environments and found it to have an uncharacteristically high relative abundances of Alphaproteobacteria, compared to the strictly marine fish they have sampled. A similar excess of Alphaproteobacteria was found in wild *Salmo salar* [18] *and Salvelinus fontinalis* [17], anadromous salmonid species. Amazon River fish were also found to have high Alphaproteobacteria under certain physicochemical conditions [31]. In stark contrast, Gammaproteobacteria dominated the skin microbiome in wild *S. salar* fry [24], and also that of *Silurus glanis*, catfish caught in the wild [16]. Of the five instances, the catfish is the only strictly-freshwater inhabitant, but it lacks scales.

In this study, we have investigated freshwater fish and identified Alphaproteobacteria as having the highest median relative abundance, highlighting their dominance as a possible feature of some freshwater fish skin microbiomes, compared to marine fish [15,46]. Such a difference is conceivable due to consistent abiotic differences between marine and freshwater habitats, and the resulting differences in fish biology between them. However, with such few and methodologically different studies of freshwater fish, and the exceptions that exist among them, this hypothesis requires further study.

### Site related factors shaping the fish skin microbiome

Based on alpha and beta diversity analyses, the raw swab communities of some of the sites are clearly different from the water communities in the same location, particularly when water temperature is low. This may form the impression that the raw swab communities properly represent the skin microbiome in fish. However, our results show that this may be misleading and the water background contamination should be formally addressed. Water measurements, especially the temperature, may seem to govern alpha and beta diversity of fish skin communities. The temperature and conductivity may be perceived as overwhelmingly strong effects on beta diversity in particular (R^2^ = 0.75 and 0.66 for the first PCoA axis). However, when the water background contamination is addressed, these effects are lost completely for alpha diversity, and become much weaker, in the case of beta diversity, where the percent dissolved oxygen and temperature emerge as two weak factors (R^2^ = 0.18 for the first axis and R^2^ = 0.15 for the second axis, for oxygen and temperature, respectively).

Statistical tests of group effects on the alpha and beta diversity support this finding. Alpha diversity differences among streams are completely lost following the elimination of background water contamination, revealing a constant alpha diversity in fish skin among streams in the system. Beta diversity significantly differs among most pairs of streams for the raw and corrected communities, but the ASVs causing these differences change with the elimination of background contamination. *Cetobacterium*, a skin microbiome salinity bioindicator [25], emerges as the main source of variation among sites, but this effect is lost following the treatment of background noise, and Phycisphaeraceae and *Anaerobacillus* become the main varying component. Both Phycisphaeraceae [43] and *Anaerobacillus* are facultative or strict anaerobes and accordingly they change with the dissolved oxygen levels.

To summarize, in the studied area, the alpha diversity of fish skin microbiomes is governed by limiting factors set by the skin mucus and are independent from the abiotic condition differences among sites. Beta diversity seems to be sensitive mainly to dissolved oxygen levels in the water, bearing in mind that dissolved oxygen measurements are missing from the H stream. This finding is consistent with the results obtained by Sylvain et al. [31] who found dissolved oxygen to be a stronger water property than the temperature, salinity and pH in shaping the skin microbiome composition in two Amazon River species. Sylvain et al. [31] identified even stronger chemical properties, which we have not accounted for in this study.

### Fish taxonomy effects on fish skin microbiomes

Alpha diversity differences between spatially overlapping fish taxa significantly emerge only when the water background contamination is addressed, highlighting once again the importance of this analytic procedure. The *O. insignis* skin microbiome has a higher alpha diversity than co-occurring species. Beta diversity differences between *O. insignis* and co-occurring species is also detected, and the statistical significance of these comparisons persist with the elimination of background contamination. We do not have the scope to determine the source of this difference, with fish phylogeny or niche among the possible sources, *O. insignis* being strictly bentic and *C. damascina*, the most common cyprinid, strictly pelagic.

### Anthropogenic eutrophication promotes skin dysbiosis

We have *a-priori* defined three sites as interrupted, in which human activity produces excess nutrients that are released into the water. These include a settling pool feeding into the Hermon Stream (site H.0.6), a site in the Snir Stream downstream a fish-farm outlet (S.3) and a fish-pond outlet on the Jordan River (J.1). The relative abundances of two phyla, Proteobacteria and Bacteroidetes, strongly covary with these levels of interrupted and uninterrupted sites, where Bacteroidetes relative abundances increase at the expense of Proteobacteria, at the interrupted sites. The relationship between these two phyla is a hallmark of dysbiosis and reduced fish health, in the skin microbiome [44]. An additional site, which we have not *a-priori* identified as interrupted, also presented elevated Bacteroidetes relative abundances. This site is shown in Fig. S2, to be a very small water body, which is likely to be easily enriched by runoff. As Fig. 6B shows, the treatment of background noise is crucial to distinguish an increase of Bacteroidetes in the water from real skin dysbiosis. This result supports the finding of Legrand et al. [44] as a useful ecological bioindicator for monitoring wild environments. Further, it is in line with the notion that sporadic pollution events of aquatic environments cannot always be detected by bulk water monitoring strategies, while biofilms do capture such events and bear testament to them [47].

### Predicted skin microbiome function changes with the consideration of background noise

To predict the implications of background noise treatment for functional inference, we compared the KEGG pathway composition between the raw and corrected predicted metagenomes. We have found that the removal of variants that equally occur in the skin and water microbiomes, fundamentally changes the variety of potential pathways in the metagenome.The largest change between the raw and corrected microbial skin communities was in KEGG pathways related to the biosynthesis of secondary metabolites, microbial metabolism in diverse environments and the biosynthesis of antibiotics.These categories are fundamental to the way bacteria interact with their environment and a metagenomic analysis of this sort would be more accurate, taking background noise effects on the microbiome composition into consideration.

## Conclusion

In this study we highlight the importance of a formal consideration of water background noise in fish skin microbiomes, when studying heterogeneous inland systems, in which fish species and environmental conditions covary. In the northern Jordan River system, north and south of the Sea of Galilee, we identify a consistent alpha diversity among sites, indicating that the limiting factors of alpha diversity in the skin microbiome are set by the mucus itself, and not by water properties. We further identify the dissolved oxygen to play a role in governing the community composition on the skin, in accordance with a previous research. Finally, we find the ratio of Proteobacteria and Bacteroidetes in the skin microbiome, a useful and informative biomarker for freshwater habitat monitoring.

## Methods

### Study area, sampling procedure and fish identification

Samples were collected between August and October 2017 from 17 sites in the Northern Jordan River water system, nine of the sites representing the upper reaches tributaries Hermon and Snir, five sites representing the northern Jordan River itself, and three additional sites from the Springs Valley Jordan River tributaries (Fig. 1, Table S1). Fish collection was commissioned by the Israeli Nature and Park Authority (NPA), as a part of their monitoring program, under permit 2017/41719. Fish were collected using either an electroshocker or a seine and placed in multiple large containers to avoid contact among individuals. The fish were classified on site, swabbed along the lateral line using a sterile swab, and released immediately. Fish species were identified according to the following criteria: *Oxynoemacheilus insignis* (Heckel, 1843) is the only loach in the system. *Astatotilapia flaviijosephi* (Lortet, 1883) is the only haplochromine in the system, and it is therefore the only cichlid with egg-shaped marks on its anal fin. *Coptodon zillii* (Gervais, 1848) is a cichlid with a dotted tail fin and 8-9 protrusions per gill raker. *Sarotherodon galilaeus* (Linnaeus, 1758) is a cichlid with a clear convex tail fin and over 13 protrusions per gill raker and a black mark on the operculum. *Oreochromis* hybrids are cichlids with striped tail fins and over 17 protrusions per gill raker. *Gambusia affinis* (Baird & Girard, 1853) is the only killifish in the study area and identifiable by its size (< 5 cm) and superior mouth. *Mugilidae* individuals escaped from fish farms in the regions (The Jordan River system has no marine outlet) and where identified by their general mugilid form. Within Cyprinidae, *Carasobarbus canis* (Valenciennes, 1842) and *Barbus longiceps* (Valenciennes, 1842) each have two pairs of barbels, *B. longiceps* with an elongated head and over 50 scales along its lateral line. *C. canis* is distinguishable by its short head and very large scales, less than 40 along the lateral line. *Capoeta damascina* (Valenciennes, 1842) has one pair of barbels and very small scales, over 70 along its lateral line. *Garra nana* (Heckel, 1843) and *Garra jordanica* Geiger & Freyhof, 2014 have small, barely visible barbels. *G. jordanica* has a suction cup and *G. nana* a fold, under the lower lip. *Acanthobrama lissneri* Tortonese, 1952, has elongated and compressed body, with deeply forked tail and up to 12 cm adult total length, and *Pseudophoxinus kervillei* (Pellegrin, 1911) is of similar size, has circular body section and a forked tail with a black stain at the base.

In addition to swab samples, in each sampling site, 2 liter of water were filtered using a sterile mixed cellulose esters 0.45 µm pore size filter. The swabs and filters were kept at in ice on site and transferred to a −80 °C until further processing. The water temperature, conductivity, pH and percent dissolved oxygen were measured at each site using a YSI ProPlus with a Quatro Cable multiparameter cable, with the exception of dissolved oxygen at Hermon Stream (H) sites and pH at two Jordan River (J) sites.

### 16S rRNA library preparation

DNA was extracted from the swabs and filters using the DNeasy PowerSoil and PowerWater DNA extraction kits (Qiagen) respectively, following the manufacturer’s instructions. Metabarcoding libraries were prepared using a two step PCR protocol, in which the first PCR reaction is designed to amplify the genetic marker along with artificial overhang sequences and the second PCR reaction is designed to attach sample specific barcode sequences and Illumina flow cell adapters. The forward and reverse PCR primers in the first reaction were ‘5-tcgtcggcagcgtcagatgtgtataagagacagCCTACG GGNGGCWGCAG-’3 and ‘5-gtctcgtgggctcggaga tgtgtataagagacagGACTACHVGGGTATCTAATC C-’3 respectively, including the target specific primers for the V3-V4 region [48] with overhangs in lowercase. For the second PCR reaction, the forward and reverse primers were ‘5-AATGATACGGCGACCACCGAGATCTACACtcgtcggcagcgtcagatgtgtataagagacag-’3 and ‘5-CAAGCAGAAGACGGCATACGAGATXXXXXXgt ctcgtgggctcgg-’3’, with Illumina adapters (uppercase), overhang complementary sequences (lowercase), and sample specific DNA barcodes (‘X’ sequence). The PCR reactions were carried out in triplicate, with the KAPA HiFi HotStart ReadyMix PCR Kit (KAPA biosystems), in a volume of 25 µl, including 1 µl of DNA template and following the manufacturer’s instructions. The first PCR reaction started with a denaturation step of 3 minutes at 95 °C, followed by 35 cycles of 20 seconds denaturation at 98 °C, 15 seconds of annealing at 55 °C and 7 seconds polymerization at 72 °C. The reaction was finalized with another one minute long polymerization step. The second PCR reaction was carried out in a volume of 25 µl as well, but with 10 µl of the PCR1 product as DNA template. It started with a denaturation step of 3 minutes at 95 °C, followed by 8 cycles of 20 seconds denaturation at 98 °C, 15 seconds of annealing at 55 °C and 7 seconds polymerization at 72 °C. The second PCR reaction was also finalized with another one minute long polymerization step. The first and second PCR reaction products were purified using AMPure XP PCR product cleanup and size selection kit (Beckman Coulter), following the manufacturer’s instructions, and sequenced on an Illumina MiSeq to produce 250 base-pair paired-end sequence reads. The sequencing was carried out by the genomics applications laboratory at the faculty of medicine, Hebrew University. The raw sequence data is archived in NCBI under BioProject PRJNA560003 (Temporary reviewer’s link: https://bit.ly/2YWRTvC).

### Amplicon sequence variance, taxonomy assignment, and background noise treatment

The bioinformatics analysis is provided on GitHub, at https://git.io/fjFZo (DOI: 10.5281/zenodo.3373312) as a jupyter notebook (https://bit.ly/2z7teFm), coupled with raw data, intermediate and output files. Sequence data trimming, amplicon sequence variant (ASV) prediction and taxonomic identification were carried out in Trimmomatic 0.39 [49] (https://bit.ly/2Hcv6AZ) and DADA2 1.12 [50]. The naive bayesian classifier used to predict taxonomic identities was trained with data from the SILVA SSU-rRNA database version 132 [51] (https://bit.ly/2OZXrkl). The resulting ASV biom table was filtered with QIIME2 2019.4 [52] to exclude ASVs assignable to eukaryotes or eukaryotic organelles, and include ones with at least 100 copies in at least two samples (https://bit.ly/30euZwh).Following alpha rarefaction analysis (https://bit.ly/2NeetsA), the ASV biom table was further filtered to exclude samples with less than 1000 sequences. A subset of the ASV biom table was created to represent the skin microbiome without ASVs that are likely to belong strictly to the water (https://bit.ly/2Z4vXhp). In this subset, we included ASVs that were unique to the swab samples, or that had a significantly higher relative abundance in the swab samples than in water samples, based on Benjamini-Hochberg corrected [33] Mann-Whitney U-test [34] (https://bit.ly/2HbEyEP). To carry out this test we used SciPy 1.2 [53] and StatsModels 0.10 [54]. The original and corrected probability values are denoted “p-value” and “q-value”, respectively. This process is regarded as “background noise treatment”, and the subset as the “corrected skin community” throughout the text. To study the taxonomic composition of the samples (https://bit.ly/2TMkEFl) and the relationship between Proteobacteria and Bacteroidetes in the different sampling sites (https://bit.ly/33GUgkL), we collapsed the ASV biom table to taxonomic tables (https://bit.ly/2z9Qlic) using QIIME2 2019.4 [52].

### Biodiversity analyses

To study the factors shaping alpha diversity, we computed Faith phylogenetic diversity (Faith PD) indices [35] for each sample, and tested the global and pairwise effect of stream and fish family levels, using the Kruskal-Wallis test [37] in QIIME2 2019.4 [52] (https://bit.ly/2OZPfR2). Faith PD depends on the number of ASVs in the sample, their pairwise phylogenetic distances and their relative abundances. We further tested the correlation of Faith PD values with the water measurements using SciPy 1.2 [53]. This was carried out for both the raw swab communities and the corrected skin communities (https://bit.ly/2z6v217).

To study the factors shaping beta diversity, we produced unweighted-UNIFRAC matrices [36] which were used for principal coordinates analysis (PCoA) [39,40], biplots [40], and PERMANOVA tests [38] in QIIME2 2019.4 [52] (https://bit.ly/2OZPfR2). The factors considered were the stream of origin, and the family or tribe of the host fish. For the latter, we carried the analyses per basin, to increase the geographic overlap of the fish species. This procedure was carried out for both the raw swab communities and corrected skin communities. We further tested the correlation of the water measurements with the values along the first and second PCoA axes, in order to explain these axes, using SciPy 1.2 [53] (https://bit.ly/2KV9Vod). Finally we executed ANCOM tests [41] to identify the bacterial ASVs explaining the group separation between significantly different fish families (https://bit.ly/2KH1M7O).

### Functional implications of background noise treatments in swab samples

To predict the differences in relative abundances of metabolic pathways between the raw and treated swab communities, we predicted their metagenomes and abundances of KEGG ENZYME terms [55], using PICRUSt 2.1.4-b [45] (https://bit.ly/2ZcYSf4). ENZYME term abundances were converted to relative abundances using pandas 0.42 [56] and the differences between the raw and corrected samples were tested with Benjamini-Hochberg corrected [33] Wilcoxon tests [57] in SciPy 1.2 [53] and StatsModels 0.10 [54]. The original and corrected probability values are denoted “p-value” and “q-value”, respectively. The KEGG PATHWAY categories of each significantly different entry were retrieved with Biopython’s REST KEGG API [58].

## Availability of data and materials

The datasets generated and analyzed during the current study are available in the National Center for Biotechnology Information (NCBI) BioProject repository under the accession number PRJNA560003. Data and script are archived as a GitHub release (https://git.io/fjFZo, DOI: 10.5281/zenodo.3370515)

## Acknowledgements

We are grateful to Dr. Dana Milshtein from Israel Nature and Parks Authority for her assistance and support.

## Funding

This research was funded by ICA in Israel, grant 03-16-06a.

## Contributions

AS and YK designed the study and collected the samples. YK identified the fish. RSA carried out the lab work. TY, AS and YA analyzed the data. TY, AS and YK wrote the manuscript. All authors read and approved the final manuscript.

## Supplementary material legends

See on FigShare https://bit.ly/2z8mqas

**Figure S1**: Alpha rarefaction curves for each fish family or tribe, denoted by the color legend. The x-axis is the size of sequence reads subsample and the y-axis is the shannon diversity in the subsample. 10.6084/m9.figshare.9642797

**Figure S2**: Site T.2 - Tel Saharonim, Springs Valley, southern basin of the study area. 10.6084/ m9.figshare.9642800

**Table S1**: The number of fish swabs collected from each fish species in each site. Sites are sorted according to stream and basin. Fish species are sorted by family or tribe. Site codes correspond with Fig. 1. N: Northern basin, north of the Sea of Galilee. S: Southern basin, south of the Sea of Galilee. IH: Exotic haplochromine. OH: *Oreochromis* hybrid. 10.6084/m9.figshare.9642824

**Table S2**: Pairwise Kruskal-Wallis tests for alpha diversity differences among streams, for raw swab bacterial communities and corrected skin bacterial communities. 10.6084/m9.figshare.9642827

**Table S3**: Pairwise Kruskal-Wallis tests for alpha diversity differences among fish families or tribes, for raw swab bacterial communities and corrected skin bacterial communities. 10.6084/m9.figshare.9642833

**Table S4**: Pairwise PERMANOVA tests for beta diversity differences among streams, for raw swab bacterial communities and corrected skin bacterial communities. 10.6084/m9.figshare.9642836

**Table S5**: Pairwise PERMANOVA tests for beta diversity differences among fish families or tribes, for raw swab bacterial communities and corrected skin bacterial communities. 10.6084/m9.figshare.9642839

## Notes

#### Summary of Updates

A few typos were corrected

https://bit.ly/2z8mqas

## References

1. Raj VS, Fournier G, Rakus K, Ronsmans M, Ouyang P, Michel B, et al. Skin mucus of *Cyprinus carpio* inhibits cyprinid herpesvirus 3 binding to epidermal cells. Vet Res. 2011;42:92.

2. Merrifield DL, Rodiles A. 10 - The fish microbiome and its interactions with mucosal tissues. In: Beck BH, Peatman E, editors. Mucosal Health in Aquaculture. San Diego: Academic Press; 2015. p. 273–95.

3. Reverter M, Tapissier-Bontemps N, Lecchini D, Banaigs B, Sasal P. Biological and ecological roles of external fish mucus: a review. Fish Sahul. 2018;3:41.

4. Ángeles Esteban M, Cerezuela R. 4 - Fish mucosal immunity: skin. In: Beck BH, Peatman E, editors. Mucosal Health in Aquaculture. San Diego: Academic Press; 2015. p. 67–92.

5. Guardiola FA, Cuesta A, Abellán E, Meseguer J, Esteban MA. Comparative analysis of the humoral immunity of skin mucus from several marine teleost fish. Fish Shellfish Immunol. 2014;40:24–31.

6. Ringø E, Holzapfel W. Identification and characterization of *Carnobacteria* associated with the gills of Atlantic salmon (*Salmo salar* L.). Syst Appl Microbiol. 2000;23:523–7.

7. Olsson JC, Westerdahl A, Conway PL, Kjelleberg S. Intestinal colonization potential of turbot (*Scophthalmus maximus*) and dab (*Limanda limanda*) associated bacteria with inhibitory effects against *Vibrio anguillarum*. Appl Environ Microbiol. 1992;58:551–6.

8. Ringø E, Olsen RE. The effect of diet on aerobic bacterial flora associated with intestine of Arctic charr (*Salvelinus alpinus* L.). J Appl Microbiol. 1999;86:22–8.

9. Olafsen JA. Interactions between fish larvae and bacteria in marine aquaculture. Aquaculture. 2001;200:223–47.

10. Balcázar JL, Vendrell D, de Blas I, Ruiz-Zarzuela I, Gironés O, Múzquiz JL. In vitro competitive adhesion and production of antagonistic compounds by lactic acid bacteria against fish pathogens. Vet Microbiol. 2007;122:373–80.

11. Pérez-Sánchez T, Balcázar JL, García Y, Halaihel N, Vendrell D, de Blas I, et al. Identification and characterization of lactic acid bacteria isolated from rainbow trout, *Oncorhynchus mykiss* (Walbaum), with inhibitory activity against *Lactococcus garvieae*. J Fish Dis. 2011;34:499–507.

12. Llewellyn MS, Boutin S, Hoseinifar SH, Derome N. Teleost microbiomes: the state of the art in their characterization, manipulation and importance in aquaculture and fisheries. Front Microbiol. 2014;5:207.

13. Brumlow CE, Luna RA, Hollister EB, Gomez JA, Burcham LA, Cowdrey MB, et al. Biochemical but not compositional recovery of skin mucosal microbiome communities after disruption. Infect Drug Resist. 2019;12:399–416.

14. Larsen A, Tao Z, Bullard SA, Arias CR. Diversity of the skin microbiota of fishes: evidence for host species specificity. FEMS Microbiol Ecol. 2013;85:483–94.

15. Chiarello M, Auguet J-C, Bettarel Y, Bouvier C, Claverie T, Graham NAJ, et al. Skin microbiome of coral reef fish is highly variable and driven by host phylogeny and diet. Microbiome. 2018;6:147.

16. Chiarello M, Paz-Vinas I, Veyssière C, Santoul F, Loot G, Ferriol J, et al. Environmental conditions and neutral processes shape the skin microbiome of European catfish (*Silurus glanis*) populations of Southwestern France. Environ Microbiol Rep. 2019;11:605–14.

17. Boutin S, Sauvage C, Bernatchez L, Audet C, Derome N. Inter individual variations of the fish skin microbiota: host genetics basis of mutualism? PLoS One. 2014;9:e102649.

18. Uren Webster TM, Consuegra S, Hitchings M, Garcia de Leaniz C. Interpopulation variation in the Atlantic Salmon microbiome reflects environmental and genetic diversity. Appl Environ Microbiol. 2018;84.

19. Svanevik CS, Lunestad BT. Characterisation of the microbiota of Atlantic mackerel (*Scomber scombrus*). Int J Food Microbiol. 2011;151:164–70.

20. Arias CR, Koenders K, Larsen AM. Predominant bacteria associated with red snapper from the Northern Gulf of Mexico. J Aquat Anim Health. 2013;25:281–9.

21. Boutin S, Bernatchez L, Audet C, Derôme N. Network analysis highlights complex interactions between pathogen, host and commensal microbiota. PLoS One. 2013;8:e84772.

22. Boutin S, Audet C, Derome N. Probiotic treatment by indigenous bacteria decreases mortality without disturbing the natural microbiota of *Salvelinus fontinalis*. Can J Microbiol. 2013;59:662–70.

23. Tarnecki AM, Brennan NP, Schloesser RW, Rhody NR. Shifts in the skin-associated microbiota of hatchery-reared common snook *Centropomus undecimalis* during acclimation to the wild. Microb Ecol. 2019;77:770.

24. Uren Webster TM, Rodriguez-Barreto D, Castaldo G, Gough P, Consuegra S, de Leaniz CG. Environmental plasticity and colonisation history in the Atlantic salmon microbiome: a translocation experiment. bioRxiv. 2019;564104.

25. Schmidt VT, Smith KF, Melvin DW, Amaral-Zettler LA. Community assembly of a euryhaline fish microbiome during salinity acclimation. Mol Ecol. 2015;24:2537–50.

26. Salinas I, Magadán S. Omics in fish mucosal immunity. Dev Comp Immunol. 2017;75:99–108.

27. Ross AA, Rodrigues Hoffmann A, Neufeld JD. The skin microbiome of vertebrates. Microbiome. 2019;7:79.

28. Rosado D, Pérez-Losada M, Severino R, Cable J, Xavier R. Characterization of the skin and gill microbiomes of the farmed seabass (*Dicentrarchus labrax*) and seabream (*Sparus aurata*). Aquaculture. 2019;500:57–64.

29. Bastos Gomes G, Hutson KS, Domingos JA, Infante Villamil S, Huerlimann R, Miller TL, et al. Parasitic protozoan interactions with bacterial microbiome in a tropical fish farm. Aquaculture. 2019;502:196–201.

30. Foysal MJ, Momtaz F, Robiul Kawser AQM, Chaklader MR, Siddik MAB, Lamichhane B, et al. Microbiome patterns reveal the transmission of pathogenic bacteria in hilsa fish (*Tenualosa ilisha*) marketed for human consumption in Bangladesh. J Appl Microbiol. 2019;126:1879–90.

31. Sylvain F-É, Holland A, Audet-Gilbert É, Luis Val A, Derome N. Amazon fish bacterial communities show structural convergence along widespread hydrochemical gradients. Mol Ecol. 2019;0:11–5.

32. Colston TJ, Jackson CR. Microbiome evolution along divergent branches of the vertebrate tree of life: what is known and unknown. Mol Ecol. 2016;25:3776–800.

33. Benjamini Y, Hochberg Y. Controlling the false discovery rate: a practical and powerful approach to multiple testing. J Royal Stat Soc B. 1995;57:289–300.

34. Mann HB, Whitney DR. On a test of whether one of two random variables is stochastically larger than the other. Ann Math Stat. 1947;18:50–60.

35. Faith DP. Conservation evaluation and phylogenetic diversity. Biol Conserv. 1992;61:1–

36. Lozupone C, Knight R. UniFrac: a new phylogenetic method for comparing microbial communities. Appl Environ Microbiol. 2005;71:8228–35.

37. Kruskal WH, Allen Wallis W. Use of ranks in one-criterion variance analysis. J Am Stat Assoc. 1952;47:583–621.

38. Anderson MJ. A new method for non-parametric multivariate analysis of variance. Austral Ecology. 2001;26:32–46.

39. Halko N, Martinsson P, Shkolnisky Y, Tygert M. An algorithm for the principal component analysis of large data sets. SIAM J Sci Comput. 2011;33:2580–94.

40. Legendre P, Legendre L. Numerical Ecology. third edition. London: Elsevier; 2012.

41. Mandal S, Van Treuren W, White RA, Eggesbø M, Knight R, Peddada SD. Analysis of composition of microbiomes: a novel method for studying microbial composition. Microb Ecol Health Dis. 2015;26:27663.

42. Pearson Karl, Galton Francis. VII. Note on regression and inheritance in the case of two parents. Proc R Soc Lond. Royal Society; 1895;58:240–2.

43. Fukunaga Y, Kurahashi M, Sakiyama Y, Ohuchi M, Yokota A, Harayama S. *Phycisphaera mikurensis* gen. nov., sp. nov., isolated from a marine alga, and proposal of Phycisphaeraceae fam. nov., Phycisphaerales ord. nov. and Phycisphaerae classis nov. in the phylum Planctomycetes. J Gen Appl Microbiol. 2009;55:267–75.

44. Legrand TPRA, Catalano SR, Wos-Oxley ML, Stephens F, Landos M, Bansemer MS, et al. The inner workings of the outer surface: skin and gill microbiota as indicators of changing gut health in Yellowtail Kingfish. Front Microbiol. 2017;8:2664.

45. Langille MGI, Zaneveld J, Caporaso JG, McDonald D, Knights D, Reyes JA, et al. Predictive functional profiling of microbial communities using 16S rRNA marker gene sequences. Nat Biotechnol. 2013;31:814–21.

46. Minich JJ, Petrus S, Michael JD, Michael TP, Knight R, Allen EE. Temporal, environmental, and biological drivers of the mucosal microbiome in a wild marine fish, *Scomber japonicus*. bioRxiv. 2019;721555.

47. Pu Y, Ngan WY, Yao Y, Habimana O. Could benthic biofilm analyses be used as a reliable proxy for freshwater environmental health? Environ Pollut. 2019;252:440–9.

48. Klindworth A, Pruesse E, Schweer T, Peplies J, Quast C, Horn M, et al. Evaluation of general 16S ribosomal RNA gene PCR primers for classical and next-generation sequencing-based diversity studies. Nucleic Acids Res. 2013;41:e1.

49. Bolger AM, Lohse M, Usadel B. Trimmomatic: a flexible trimmer for Illumina sequence data. Bioinformatics. 2014;30:2114–20.

50. Callahan BJ, McMurdie PJ, Rosen MJ, Han AW, Johnson AJA, Holmes SP. DADA2: High-resolution sample inference from Illumina amplicon data. Nat Methods. 2016;13:581–3.

51. Quast C, Pruesse E, Yilmaz P, Gerken J, Schweer T, Yarza P, et al. The SILVA ribosomal RNA gene database project: improved data processing and web-based tools. Nucleic Acids Res. 2013;41:D590–6.

52. Bolyen E, Rideout JR, Dillon MR, Bokulich NA, Abnet CC, Al-Ghalith GA, et al. Reproducible, interactive, scalable and extensible microbiome data science using QIIME 2. Nat Biotechnol. 2019;37:852–7.

53. Oliphant TE. Python for Scientific Computing. Computing in Science Engineering. 2007;9:10–20.

54. Seabold S, Perktold J. Statsmodels: Econometric and statistical modeling with python. 9th Python in Science Conference. 2010.

55. Kanehisa M, Sato Y, Furumichi M, Morishima K, Tanabe M. New approach for understanding genome variations in KEGG. Nucleic Acids Res. 2019;47:D590–5.

56. McKinney W. Data structures for statistical computing in python. In: van der Walt S, Millman J, editors. Proceedings of the 9th Python in Science Conference. Austin, TX; 2010. p. 51–6.

57. Wilcoxon F. Individual comparisons by ranking methods. Biometrics Bulletin. 1945;1:80–3.

58. Cock PJA, Antao T, Chang JT, Chapman BA, Cox CJ, Dalke A, et al. Biopython: freely available Python tools for computational molecular biology and bioinformatics. Bioinformatics. 2009;25:1422–3.

